# Inducible Bronchus-Associated Lymphoid Tissue in SARS-CoV-2 Infected Rhesus Macaques

**DOI:** 10.1101/2022.07.12.499813

**Authors:** Zhong-Min Ma, Katherine J. Olstad, Koen K A Van Rompay, Smita S. Iyer, Christopher J. Miller, Rachel J Reader

## Abstract

Pulmonary immunity against SARS-CoV-2 infection has not been well studied. This study investigated the distribution of immune cells int the lungs of 8 rhesus macaques experimentally infected with SARS-CoV-2, and euthanized 11-14 days later. Using immunohistochemistry, inducible bronchus-associated lymphoid tissue was found in all animals. The inducible bronchus-associated lymphoid tissues were composed of B cells, T cells, and follicular dendritic cells with evidence of lymphocyte priming and differentiation. This suggests local immunity plays an important role in the SARS-CoV-2 infection. Further study of local immunity in the lung would benefit our understanding of SARS-CoV-2 pathogenesis and could lead to new interventions to control the SARS-CoV-2 infection and disease.

## Introduction

According to the WHO, to date SARS CoV-2 has infected more than 554 million people and caused more than 6,352,851 mortalities worldwide (https://covid19.who.int: accessed July 22nd 2022). While vaccine have been relatively successful, a big challenge is the continuous emergence of variants of concern with unpredictable virulence and variable impact on vaccine efficacy ^1^. Above all, understanding the pathogenesis, including development of local immune responses to infection, is fundamental for developing interventions aimed at controlling the viral infection.

The classical pathology investigations, such as postmortem examination and histology address initial questions about pathogenesis of SARS CoV-2 infection. However, human materials from COVID-19 patients which can be used for the pathological study are limited. Currently, pathological knowledge about COVID-19 comes from some postmortem biopsies and limited number of autopsies^2^. All these materials represent the latest stage of the disease, and only less than 5% of infected individuals reached this stage^3–6^. Furthermore, human sample collection usually happened more than 24 hours after death which significantly limit additional assays and data quality. Therefore, animal models play a valuable role in the study of pathogeneses and prognosis of SARS-CoV-2 infection.

Several non-human primate species (NHP), including rhesus and cynomolgus macaques and African green monkeys have been shown to be useful animal models of SARS-CoV-2, as infection mimics human SARS-CoV-2 infection in many aspects of the disease^7^, including asymptomatic or mild clinical symptoms in the majority of infections, but with histological lesions consistent with mild to moderate interstitial pneumonia. ^8, 9^ Knowledge gained by studying NHP should assist in understanding the disease, testing the efficacy of vaccination and treatment, studying sequelae following SARS-CoV-2 infection, and guiding to developing strategies for defeating COVID-19.

The inducible bronchus-associated lymphoid tissue (iBALT) is a tertiary lymphoid organ that forms in the lung following inflammation or infection and located at the basal site of the bronchial epithelium and perivascular space ^10–12^. Unlike secondary lymphoid organs, such as the spleen and lymph node, the iBALT is not encapsulated and does not constitutively exist in all mammals including mice, monkeys, and humans^12–15^. It is induced by exposure to antigens under infection or inflammatory stimulation^16, 17^. IL-1 alpha, IL-23, IL-17, IL-22, CCL19, CCL21, CXCL12, and CXCL13 and their receptors such as CCR7, CXCR4, and CXCR5 play essential roles in iBALT’s formation and development^10–12, 18^. The iBALT is characterized by a cluster of B cells, T cells, and follicular dendritic cells, which organize into a unique structure resembling lymphoid follicle^18^. Although iBALT is not extensively studied, there is evidence that it plays both a protective and pathological role in pulmonary disease^10, 11^. The development of iBALT has not been reported in COVID-19.

In this study, we performed a descriptive investigation on lungs of rhesus macaques harvested from 11-14 days post SARS-CoV-2 inoculation, and utilized histopathology and immuno-histochemistry to characterize the histopathological lesions and the distribution of immune cells. Results showed that iBALT was well established in all experimental animals. They are composed of B cells, T cells, and follicular dendritic cells with evidence of lymphocyte priming and differentiation. Our results suggested that local immunity may play an essential role in the pathogenesis of COVID-19. Further study of local immunity in the lung was necessary, and would benefit to understand SARS-CoV-2 pathogenesis and could lead to new interventions to control the SARS-CoV-2 infection and disease.

## Material and Methods

### Animal information

Detailed information on the study’s experimental design was reported earlier^19, 20^. Briefly, this study included 5 males and 3 females colony-bred Indian origin rhesus macaques (*Macaca mulatta*) with ages ranging from 4 to 5 years old and weight ranging from 5.4 to 10.7 kg (median 8.6 kg). All animals were inoculated with a California isolate of SARS-CoV-2 by intratracheal, intranasal and ocular routes. Viral RNA and SARS CoV-2 antibodies were monitored to confirm the SARS-CoV-2 infection. The study was approved by the UC-Davis Institutional Animal Care and Use Committee.

### Pathology and immune-histochemistry

Animals were euthanized at 11 to 14 days after the viral inoculation. Lungs were infused gently with 10% buffered formalin within 20 minutes after being harvested from animals at necropsy. Infused lungs were further fixed in 10% buffered formalin for 3 days and then transferred into 70% ethanol and processed to make paraffin blocks. Four µm sections were used for Hematoxylin-Eosin (HE) and immunohistochemistry (IHC) stains. Resources and the working dilution of antibodies show in Table1. All first antibodies were subjected to an antigen retrieval step consisting of incubation in AR10 (Biogenex, San Ramon CA) for 2 minutes at 125°C in the Digital Decloaking Chamber (Biocare, Concord CA) which was followed by cooling to 90°C for 10 minutes, or incubation in antigen unmasking solution H3300 (Vector, Burlingame, CA) for 20 minutes at 100°C before rinsing in water (Table 1). Primary antibodies were replaced by normal rabbit IgG or mouse IgG (Invitrogen, Rockford IL) and included in each staining series as the negative control. EnVision and AEC (Dako, Santa Clara CA) were used as detection systems. Slides were counterstained with hematoxylin, dehydrated, cover-slipped, and visualized by using a bright field microscope. In the immunofluorescent stains of CD3/PD-1 and CD3/CD20/Bcl-1, Goat anti Rat Alexa Fluor 488 and Goat anti-Rabbit Alexa Fluor 568, and Goat anti Rat Alexa Fluor 488, Goat anti Rabbit Alexa Fluor 647 and Goat anti-Mouse Alexa Fluor 568 (Invitrogen, Rockford IL) were used. With appropriate filters, slides were visualized with epi-fluorescent illumination using a Zeiss Imager microscope (Carl Zeiss Inc., Thornwood, NY).

**Table 1:** Antibodies’ resources and working condition.

| **Ab name** | **Species** | **Clone** | **Vender** | **Cat#** | **Dilution** | **Antigen Retrieval Method** |
| --- | --- | --- | --- | --- | --- | --- |
| **CD3** | Rabbit |  | DAKO, Santa Clara CA | A-0452 | 1:100 | AR10 |
| **CD3** | Rat | CD3-12 | abcam, Wattham MA | Ab11089 | 1:100 | H3300 |
| **CD4** | mouse | 1F6 | abcam, Wattham MA | ab846 | 1:25 | AR10 |
| **CD8** | Rabbit |  | DBS, Pleasanton CA | RP-100 | 1:50 | AR10 |
| **CD20** | mouse | L26 | DAKO, Santa Clara CA | M0755 | 1:300 | AR10 |
| **CD20** | Rabbit |  | DAKO, Santa Clara CA | A-0452 | 1:100 | AR10 |
| **MPO** | Rabbit |  | abcam, Wattham MA | ab9535 | 1:50 | AR10 |
| **CD68** | mouse | KP1 | Thermo Fisher, Tremont CA | MS-397PO | 1:100 | AR10 |
| **CD163** | Mouse | 10D6 | Novus, Centennial CO | NB110-59935 | 1:40 | H3300 |
| **CD169** | Rabbit | SP213 | LSBio, Seattle WA | LS-C210436 | 1:100 | H3300 |
| **Caspase 3** | Rabbit | C92-605 | BD, Franklin Lakes NJ | 559565 | 1:200 | AR10 |
| **Ki67** | Rabbit |  | Neomarkers, Fremont CA | RB1510 | 1:300 | AR10 |
| **IgG** | Rabbit |  | DAKO, Santa Clara CA | A0423 | 1:1000 | AR10 |
| **Perforin** | mouse |  | Mabtech, Cincinnati, OH | Pf-16:17 | 1:100 | AR10 |
| **Granzyme B** | mouse | GZB01 | Neomarkers, Fremont CA | MS-1157-R7 | Prediluted | AR10 |
| **PD1** | Rabbit |  | Novus, Centennial CO | NBP1-88104 | 1:100 | H3300 |
| **Bcl-6** | mouse | LN22 | Biocare, Concord CA | CM 410 A | 1:100 | AR10 |
| **p55** | mouse | 55K-2 | DAKO, Santa Clara CA | M3567 | 1:200 | AR10 |

## Results

### SARS CoV-2 infected all animals

As described earlier^19, 20^, all animals were SARS CoV-2 infected. No animals showed severe clinical signs (such as acute fever, weight loss, or respiratory distress) after infection. Clinical signs were absent or mild.

### General histopathology findings

Gross examination showed the pleural surface of lungs were smooth with randomly distributed, variable-sized, and irregular-shaped dark or pale discolorations. No pleural effusion was seen. Enlargement of lymph nodes at the hilum and mediastinum was noticed in some animals. In cross-sections, small patches of the discolorations were randomly distributed in different lobes of the lungs with evidence of mild pulmonary edema. Histopathological examination of the lungs showed multifocal to locally extensive interstitial pneumonia of mild severity frequently radiating out from the terminal bronchioles and sometimes subpleural in all animals. Histology in these regions showed that alveolar septa were expanded by a few to moderate numbers of neutrophils and mononuclear cells. Alveolar spaces variably contained macrophages, a few neutrophils, and occasional cellular debris. In some areas, there was occasional type 2 pneumocyte hyperplasia. Perivascular cuffing was present to variable degrees in all animals.

### Immunohistochemistry results

#### 1. Characterization of iBALT

iBALT was found in all animals. In addition to well-developed lymphoid follicles with a germinal center associated with bronchi and bronchioles, a significant amount of the iBALT is located in the perivascular space. The iBALT was of variable size and was randomly distributed throughout the lung parenchyma (Figure 1). The major constituents of the iBALT were CD20+ B cells. The B cells were arranged in prominent follicular structures which was sometimes eccentric and sometimes had a germinal center. CD20 and Bcl6 double-positive cells, Ki67 positive cells, Caspase-3 positive cells, and CD21 follicular dendritic cells were noticed in the geminal center (Figure 1,2). CD3+ T cells, which were mostly CD4+, were scattered within the B cell zone or polarized at one edge of the iBALT. A few CD8+ cells were present throughout the iBALT. Some of the CD3+ cells were PD-1 positive (Figure 2). Some CD169+, CD163+, and CD68+ macrophages were also found in the iBALT. Most of the CD163+, and CD68+ cells were at the edge of the B cell zone, while CD169 + cells were found at the center of the germinal center. A few IgG-positive cells were found within the iBALT. Further, in the iBALT that appeared as more homogeneous aggregates without classical B cell, T cell zone, and well-developed geminal center, relatively more Ki-67+ cells were found. This suggests that the classical iBALT originated from small aggregations of immune cells. Thus, an adaptive humoral immune response was taking place in the lungs of monkeys infected with SARS CoV-2.

**Figure 1:**
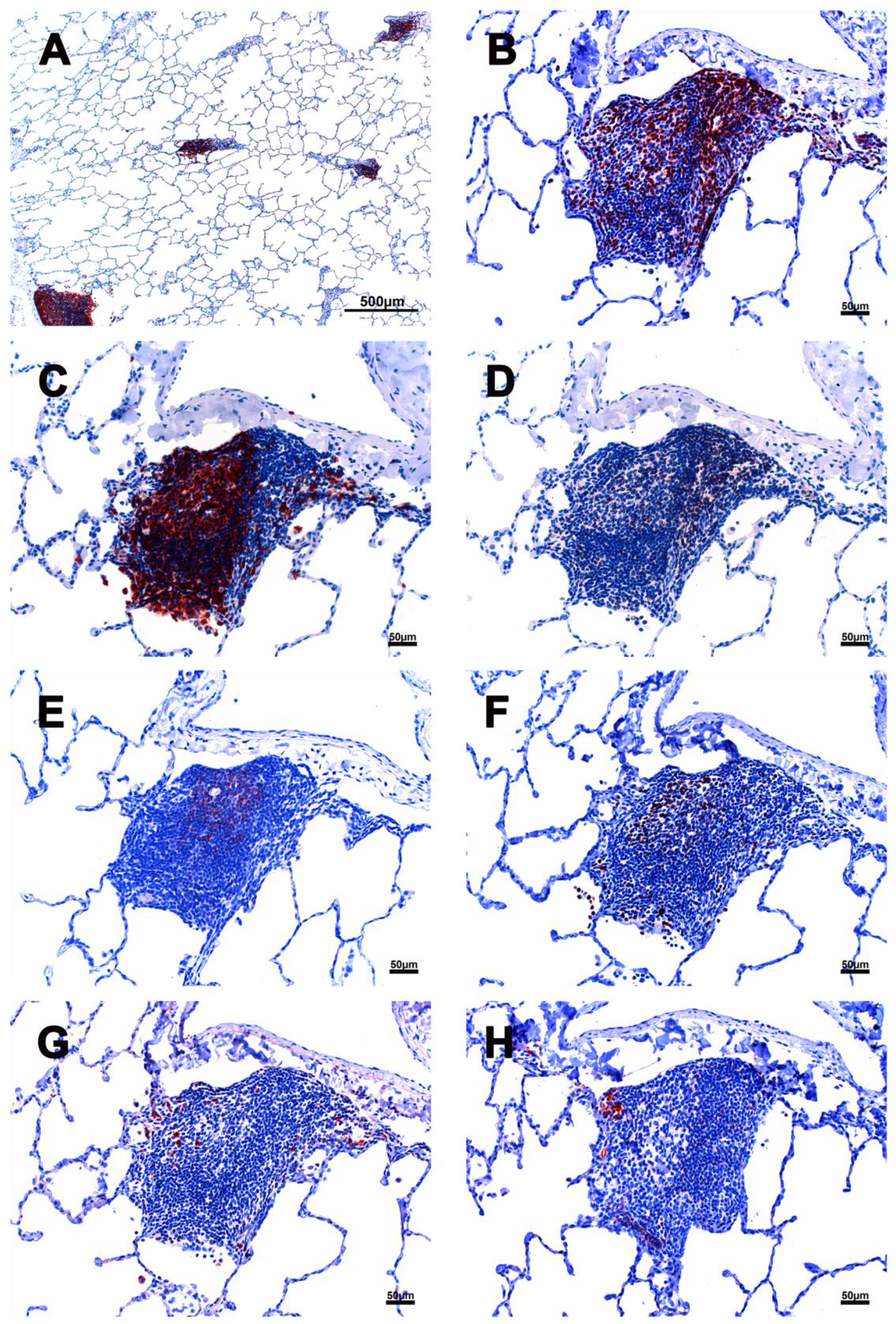
Cellular composition of iBALT of rhesus macaques which infected with SARS CoV-2. A: Multiple iBALTs identified by CD20 IHC stain. The size of the iBALTs was variable. B: CD3 positive cells aggregated along the periphery and scattered throughout the entirety of the iBALT. C: CD20 positive cells mimic a germinal center. D: CD4 positive cells. E: CD21 positive cells located in the germinal center with a dendritic structure. F: Ki67 positive cells were seen in the germinal center. G: IgG positive cells were found within the iBALT. H. P55 positive cells were found in the iBALT and identified some endothelial cells which formed well-defined small vessel structures. Scale bar on panel A was 500µm. Scale bars on panel B, C, D, E, F, G, and H were 50µm.

**Figure 2:**
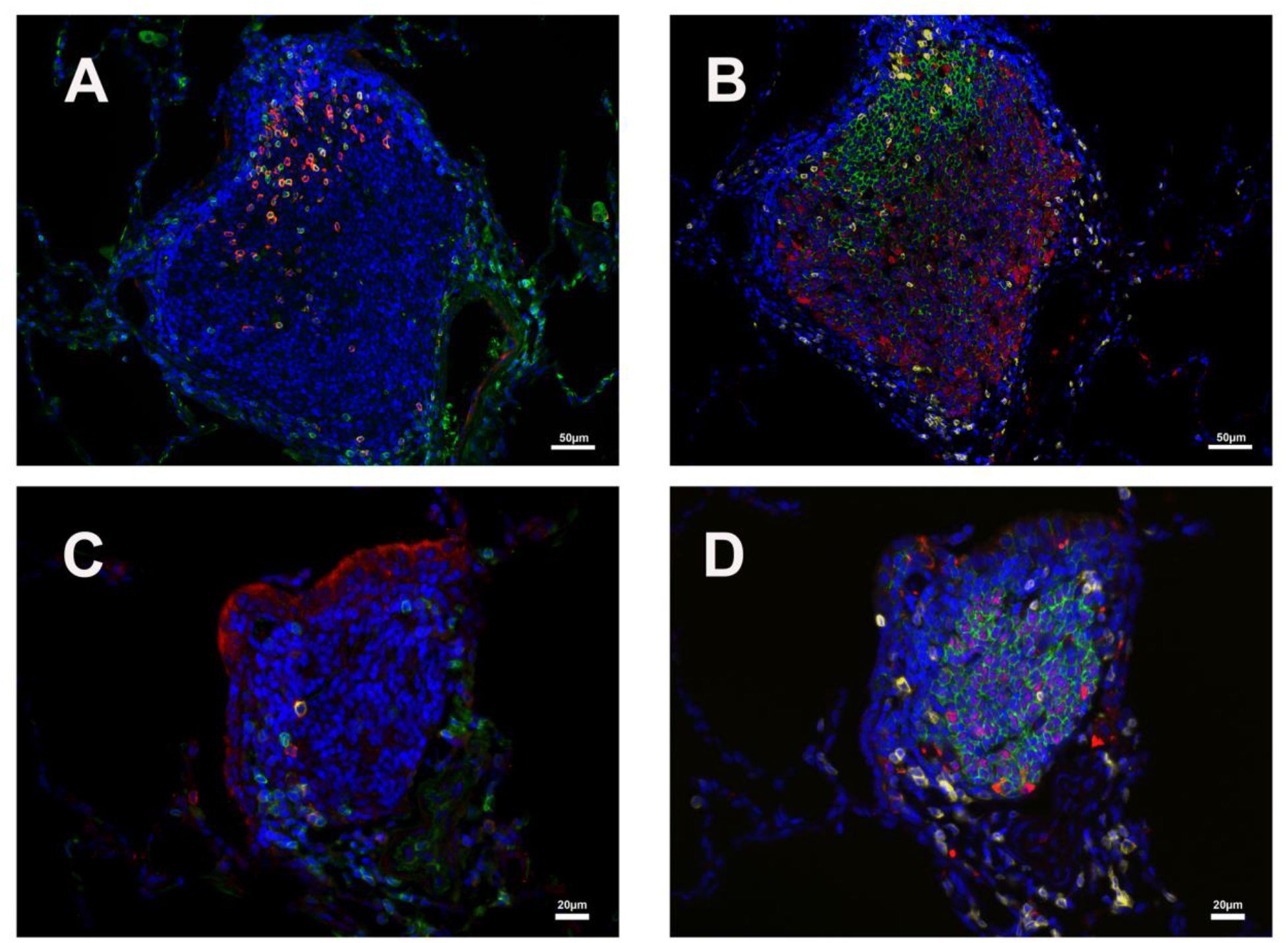
CD3/PD1 and CD20/Bcl6 double-positive cells in two iBALTs. A&C: CD3 (green) and PD1 (red) double-positive cell in the geminal center of iBLATs of different sizes. B&D: CD20 (green) and Bcl-6 (red) double-positive cell in B cell zone of iBLATs of different sizes. CD3 (yellow) positive cells are also shown. Scale bar on panel A and B were 50µm. Scale bars on panel C and D were 20µm.

The P55 stain identified some endothelial cells which formed well-defined small vessel structures inside the iBALT. Some CD21+ cells were found around small blood vessels where lymphocytes existed. Perforin and granzyme B + cells were occasionally seen in iBALT.

### 2. Immune cells beyond the iBALT areas of the lung

The distribution of immune cells in other parts of the lung, beyond the iBALT areas, was also investigated.

Myeloperoxidase (MPO) positive neutrophils were found in blood vessels, alveolar walls, alveoli, and around small blood vessels. There were more MPO + cells in the exudate within alveoli and focal areas with interstitial pneumonia. In a few cases, neutrophils infiltrated the depth of the bronchial wall to appear in the bronchial lumen. There were more MPO+ cells than CD3+ T cells, especially in the areas with interstitial pneumonia. CD3+ cells were evenly distributed in the lung with no significant difference in the number of these cells regardless of pathology. Most of the CD3+ cells were found around small blood vessels and formed aggregations in the space between blood vessels and alveoli, which colocalized with the perivascular cuffing on HE slides. Only a few CD3 + cells are located in the alveolar walls. In contrast to CD3+ T cells, CD20+ B cells were not only the primary element of iBALT but also frequently found inside alveoli and within alveolar walls. CD4 and especially CD8 staining was weak. As described previously, most of the CD4+ and CD8 + cells are found within iBALT and even fewer of them were found scattered in the alveolar walls.

Perforin + cells were found in all animals, and most of them were macrophages, located in alveoli. There were relatively more perforin+ cells within the areas with pathology. Granzyme B identified different cell populations than the perforin stain. Most granzyme B positive cells were found in the alveolar walls. The size of the granzyme B positive cells is much smaller than that of perforin positive cells.

CD68, CD163, and CD169 positive macrophages were examined in the current study. There were numerous CD68 positive macrophages in the lumen of bronchi and inside the alveoli. Multinuclear CD68 cells were found in the alveoli. Adjacent to the lymphoid aggregations, there were some macrophages with weak CD68 positive. Foreign materials were found in their cytoplasm. Those cells were also CD163, Caspase-3, and perforin positive. CD163+ and CD169 + cells, which were a little larger in size than CD68+ cells, shared a similar distribution pattern with CD68+ cells beyond the iBALT areas of the lung. However, CD169 staining was very weak in macrophages within the lumen of bronchi and within alveoli. and coincided with foreign materials in the cytoplasm. In addition to solid positive CD169+ cells in the center of the germinal center, more CD169+ were found around small blood vessels. Some spindle-shaped CD169+ cells also existed around larger blood vessels.

## Discussion

In humans, nonhuman primates, and mice, bronchus-associated lymphoid tissue (BALT) is a tertiary lymphoid organ that does not exist at birth^13, 14, 21^. Classical BALT resembles a variably sized secondary lymph follicle in a lymph node composed of many B cells in the center with surrounding T cells. Unlike most secondary lymphoid organs such as lymph nodes and spleen, which develop embryonically without microbial stimulation or antigen exposure, BALT develops after hosts have been exposed to microbial agents or experienced pulmonary disease^13^. BALT has been renamed as inducible bronchus-associated lymphoid tissue, iBALT, to reflect this property^12, 18^. In humans, the frequency of iBALT is relatively higher in children compared to people over 20 years old^22, 23^. Rabdall, T.D. et al. reported that the frequency of BALT is less than 5 per 10 histology sections in groups of young adult rhesus monkeys from the same facility as this study^12^. As the name BALT indicates, early investigators considered that BALT was located within the lamina propria of bronchi and belonged to the category of mucosal-associated lymph tissue (MALT). However, recent studies showed that iBALT can also be found in the perivascular space, which may be easily ignored when the cluster of immune cells is small, or the examined lung section is not processed correctly^15, 24, 25^. This study showed iBALT existed in both perivascular space and within bronchial wall, and most of it was located in perivascular space. So, we suggest change the name of iBALT into inducible pulmonary associated lymph tissue (iPALT) to emphasize this system was a local immunity of lung, not only associated with bronchi.

Studies show iBALT is not simply an accumulation of immune cells. Instead, these newly formed lymphoid tissues make a new micro-environment that recruits immune cells and organizes them into a tertiary lymphoid tissue that supports lymphocyte priming and differentiation^12, 18, 26, 27^. It is controversial whether or not iBALT plays a protective or pathogenic role when the host is infected by specific pathogens or under some disease conditions. However, some studies show iBALT plays a vital role in viral clearance and reducing inflammatory responses after a viral infection such as flu.Furthermore, iBALT induced by one pathogen can affect the response to a second, unrelated pathogen^28, 29^. Using a mice model, people have successfully induced immunity against two different influenza viruses, a mouse-adapted SARS-coronavirus or mouse pneumovirus infection, by inhaling Protein Cage Nanoparticle (PCN). Mice treated with the PCN show a significantly high survival rate, quick viral clearance, and accelerated viral-specific antibody production^29^. In addition, once iBALT is formed, memory T cells can be maintained, and iBALT can persist for months on site ^25^. Thus, agent that establish iBALT can potentially have a benefit in controlling future infections.

Similar to influenza virus infected mice^18^, our study shows that iBALTs are well developed after SARS-CoV-2 infection. They are scattered throughout the parenchyma of the lung, where they are not only found in the upper bronchi but also vascular space without associated bronchioles in the peripheral lung. The size and shape of iBALT varies as may its structure. Some iBALTs that possess well-developed lymph follicles were found within the wall of the upper bronchi. When secondary follicles were present, they consisted primarily of centrally located B cells that were predominantly CD20+ and less often Bcl-6 positive. T-lymphocytes forming rim around the germinal center and less often scattered within B-cell zones were predominantly CD3+and CD4+ and rarely PD-1 positive. There were Ki67, Caspase-3, and IgG-positive cells in the follicle. Dendritic cells and macrophages were also noticed within the follicle. This evidence supports a role for iBALT in lymphocyte priming and differentiation. Similar as previous study that germinal centers presented in the mediastinal lymph node following CoV-2 infection^19^, the germinal centers also existed in the iBLATs. Besides the iBALTs with a well-developed lymph follicle structure, there were many small size iBALTs located in the vascular space without any association with bronchioles in the peripheral lung. With H&E stains, these small iBALTs are easily interpreted as peri-vascular cuffs and even ignored, especially when the lung was not appropriately perfused. With IHC stains, these iBALTs were obvious and had a similar immunophenotype of CD20/BLC2+ B cells, CD3/PD-1/CD4 positive T cells, and few dendritic cells and macrophages, as described in the larger iBALT despite the lack of germinal centers.

Local immunity against SARS-CoV-2 infection, specifically iBALTs, has not been well characterized in humans partly because of the lack of samples, especially during the early stages of disease. We do not know what role iBALT are playing during the disease process. The majority of SARS-CoV-2 samples taken from infected humans is at time of autopsy when primary changes induced by the virus, such as iBALT, are obscured by a severe inflammatory background associated with mortality. As such the role of iBALT in the disease course is currently unknown. However, evidence shows that outcomes of SARS-CoV-2 infection vary within the different age groups of people and different species^30, 31^. To shed further light mechanisms behind those differences, it is worth exploring the role that local immunity, including iBALT, plays during SARS-CoV-2 infection, in particular what constitutes a healthy iBALT response (associated with no or mild disease) and whether iBALT is involved in triggering a fulminant inflammatory response that leads to severe disease and mortality. This paper is descriptive. However, the finding of this study suggested that further insights may lead to the development of the new interventions to prevent or treat SARS-CoV-2 infection and disease.

## Acknowledgments

The authors are grateful to Timothy D Carroll, Aaron Mark Allen, Sarah Lockwood and Elizabeth Rodriguez for their technical support. The authors are grateful to Greg Hodges for facilitating the animal experiments in CNPRC ABSL3. We are extremely grateful to the primate center staff Wilhelm Von Morgenland, Miles Christensen, David Bennet, Vanessa Bakula for animal sampling. We thank John H Morrison, Jeffrey A Roberts and the CNPRC leadership for their support in facilitating these experiments.

## Reference

[1] Markov PV, Katzourakis A, Stilianakis NI: Antigenic evolution will lead to new SARS-CoV-2 variants with unpredictable severity. Nat Rev Microbiol 2022.

[2] Mohanty SK, Satapathy A, Naidu MM, Mukhopadhyay S, Sharma S, Barton LM, Stroberg E, Duval EJ, Pradhan D, Tzankov A, Parwani AV: Severe acute respiratory syndrome coronavirus-2 (SARS-CoV-2) and coronavirus disease 19 (COVID-19) - anatomic pathology perspective on current knowledge. Diagn Pathol 2020, 15:103.

[3] Nishiura H, Kobayashi T, Miyama T, Suzuki A, Jung SM, Hayashi K, Kinoshita R, Yang Y, Yuan B, Akhmetzhanov AR, Linton NM: Estimation of the asymptomatic ratio of novel coronavirus infections (COVID-19). Int J Infect Dis 2020, 94:154–5.

[4] Menachemi N, Yiannoutsos CT, Dixon BE, Duszynski TJ, Fadel WF, Wools-Kaloustian KK, Unruh Needleman N, Box K, Caine V, Norwood C, Weaver L, Halverson PK: Population Point Prevalence of SARS-CoV-2 Infection Based on a Statewide Random Sample - Indiana, April 25-29, 2020. MMWR Morb Mortal Wkly Rep 2020, 69:960–4.

[5] Yang R, Gui X, Xiong Y: Comparison of Clinical Characteristics of Patients with Asymptomatic vs Symptomatic Coronavirus Disease 2019 in Wuhan, China. JAMA Netw Open 2020, 3:e2010182.

[6] Wu ZY, McGoogan JM: Characteristics of and Important Lessons From the Coronavirus Disease 2019 (COVID-19) Outbreak in China Summary of a Report of 72 314 Cases From the Chinese Center for Disease Control and Prevention. Jama-J Am Med Assoc 2020, 323:1239–42.

[7] Casel MAB, Rollon RG, Choi YK: Experimental Animal Models of Coronavirus Infections: Strengths and Limitations. Immune Netw 2021, 21:e12.

[8] Rockx B, Kuiken T, Herfst S, Bestebroer T, Lamers MM, Munnink BBO, de Meulder D, van Amerongen G, van den Brand J, Okba NMA, Schipper D, van Run P, Leijten L, Sikkema R, Verschoor E, Verstrepen B, Bogers W, Langermans J, Drosten C, van Vlissingen MF, Fouchier R, de Swart R, Koopmans M, Haagmans BL: Comparative pathogenesis of COVID-19, MERS, and SARS in a nonhuman primate model. Science 2020, 368:1012–+.

[9] Shan C, Yao YF, Yang XL, Zhou YW, Gao G, Peng Y, Yang L, Hu X, Xiong J, Jiang RD, Zhang HJ, Gao XX, Peng C, Min J, Chen Y, Si HR, Wu J, Zhou P, Wang YY, Wei HP, Pang W, Hu ZF, Lv LB, Zheng YT, Shi ZL, Yuan ZM: Infection with novel coronavirus (SARS-CoV-2) causes pneumonia in Rhesus macaques. Cell Res 2020, 30:670–7.

[10] Hwang JY, Randall TD, Silva-Sanchez A: Inducible Bronchus-Associated Lymphoid Tissue: Taming Inflammation in the Lung. Frontiers in Immunology 2016, 7.

[11] Marin ND, Dunlap MD, Kaushal D, Khader SA: Friend or Foe: The Protective and Pathological Roles of Inducible Bronchus-Associated Lymphoid Tissue in Pulmonary Diseases. J Immunol 2019, 202:2519–26.

[12] Randall TD: Bronchus-associated lymphoid tissue (BALT) structure and function. Adv Immunol 2010, 107:187–241.

[13] Pabst R, Gehrke I: Is the bronchus-associated lymphoid tissue (BALT) an integral structure of the lung in normal mammals, including humans? Am J Respir Cell Mol Biol 1990, 3:131–5.

[14] Pabst R, Miller LA, Schelegle E, Hyde DM: Organized lymphatic tissue (BALT) in lungs of rhesus monkeys after air pollutant exposure. Anat Rec (Hoboken) 2020, 303:2766–73.

[15] Woodland DL, Randall TD: Anatomical features of anti-viral immunity in the respiratory tract. Semin Immunol 2004, 16:163–70.

[16] Ruddle NH, Akirav EM: Secondary Lymphoid Organs: Responding to Genetic and Environmental Cues in Ontogeny and the Immune Response. J Immunol 2009, 183:2205–12.

[17] Gould SJ, Isaacson PG: Bronchus-Associated Lymphoid-Tissue (Balt) in Human Fetal and Infant Lung. J Pathol 1993, 169:229–34.

[18] Moyron-Quiroz JE, Rangel-Moreno J, Kusser K, Hartson L, Sprague F, Goodrich S, Woodland DL, Lund FE, Randall TD: Role of inducible bronchus associated lymphoid tissue (iBALT) in respiratory immunity. Nat Med 2004, 10:927–34.

[19] Shaan Lakshmanappa Y, Elizaldi SR, Roh JW, Schmidt BA, Carroll TD, Weaver KD, Smith JC, Verma A, Deere JD, Dutra J, Stone M, Franz S, Sammak RL, Olstad KJ, Rachel Reader J, Ma ZM, Nguyen NK, Watanabe J, Usachenko J, Immareddy R, Yee JL, Weiskopf D, Sette A, Hartigan-O’Connor D, McSorley SJ, Morrison JH, Tran NK, Simmons G, Busch MP, Kozlowski PA, Van Rompay KKA, Miller CJ, Iyer SS: SARS-CoV-2 induces robust germinal center CD4 T follicular helper cell responses in rhesus macaques. Nat Commun 2021, 12:541.

[20] Deere JD, Carroll TD, Dutra J, Fritts L, Sammak RL, Yee JL, Olstad KJ, Reader JR, Kistler A, Kamm J, Di Germanio C, Shaan Lakshmanappa Y, Elizaldi SR, Roh JW, Simmons G, Watanabe J, Pollard RE, Usachenko J, Immareddy R, Schmidt BA, O’Connor SL, DeRisi J, Busch MP, Iyer SS, Van Rompay KKA, Hartigan-O’Connor DJ, Miller CJ: SARS-CoV-2 Infection of Rhesus Macaques Treated Early with Human COVID-19 Convalescent Plasma. Microbiol Spectr 2021, 9:e0139721.

[21] Tschernig T, Pabst R: Bronchus-associated lymphoid tissue (BALT) is not present in the normal adult lung but in different diseases. Pathobiology 2000, 68:1–8.

[22] Hiller AS, Tschernig T, Kleemann WJ, Pabst R: Bronchus-associated lymphoid tissue (BALT) and larynx-associated lymphoid tissue (LALT) are found at different frequencies in children, adolescents and adults. Scand J Immunol 1998, 47:159–62.

[23] Emery JL, Dinsdale F: Increased Incidence of Lymphoreticular Aggregates in Lungs of Children Found Unexpectedly Dead. Arch Dis Child 1974, 49:107–11.

[24] Pabst R, Tschernig T: Perivascular capillaries in the lung: an important but neglected vascular bed in immune reactions? J Allergy Clin Immunol 2002, 110:209–14.

[25] Moyron-Quiroz JE, Rangel-Moreno J, Hartson L, Kusser K, Tighe MP, Klonowski KD, Lefrancois L, Cauley LS, Harmsen AG, Lund FE, Randall TD: Persistence and responsiveness of immunologic memory in the absence of secondary lymphoid organs. Immunity 2006, 25:643–54.

[26] Moyron-Quiroz J, Rangel-Moreno J, Carragher DM, Randall TD: The function of local lymphoid tissues in pulmonary immune responses. Adv Exp Med Biol 2007, 590:55–68.

[27] Hwang JY, Randall TD, Silva-Sanchez A: Inducible Bronchus-Associated Lymphoid Tissue: Taming Inflammation in the Lung. Front Immunol 2016, 7:258.

[28] Halle S, Dujardin HC, Bakocevic N, Fleige H, Danzer H, Willenzon S, Suezer Y, Hammerling G, Garbi N, Sutter G, Worbs T, Forster R: Induced bronchus-associated lymphoid tissue serves as a general priming site for T cells and is maintained by dendritic cells. J Exp Med 2009, 206:2593–601.

[29] Wiley JA, Richert LE, Swain SD, Harmsen A, Barnard DL, Randall TD, Jutila M, Douglas T, Broomell C, Young M, Harmsen A: Inducible bronchus-associated lymphoid tissue elicited by a protein cage nanoparticle enhances protection in mice against diverse respiratory viruses. PLoS One 2009, 4:e7142.

[30] Qin S, Li R, Zheng Z, Zeng X, Wang Y, Wang X: Review of selected animal models for respiratory coronavirus infection and its application in drug research. J Med Virol 2022.

[31] Hu B, Guo H, Zhou P, Shi ZL: Characteristics of SARS-CoV-2 and COVID-19. Nat Rev Microbiol 2021, 19:141–54.

